# Enhancing Cardiac Reprogramming by Suppressing Specific C-C Chemokine Signaling Pathways

**DOI:** 10.1101/522995

**Authors:** Yijing Guo, Ienglam Lei, Shuo Tian, Wenbin Gao, Karatas Hacer, Yangbing Li, Shaomeng Wang, Liu Liu, Zhong Wang

## Abstract

Reprogramming fibroblasts into induced cardiomyocytes (iCMs) is a potentially promising strategy for heart regeneration. Yet a major challenge is the low conversion rate. To address this challenge, we screened and identified four chemicals, insulin-like growth factor-1, Mll1 inhibitor MM589, transforming growth factor-β inhibitor A83-01, and Bmi1 inhibitor PTC-209, termed as IMAP, that coordinately enhanced reprogramming efficiency.

Using α-muscle heavy chain -green fluorescent protein mouse embryo fibroblasts as the staring cell type, we observed a six-fold increase of iCM formation with IMAP treatment. IMAP stimulated higher cardiac troponin T and α-actinin expression and more sarcomere formation with up-regulation of many cardiac genes and down-regulation of fibroblast genes. Furthermore, IMAP promoted higher spontaneous beating and calcium transient activities of iCMs derived from neonatal cardiac fibroblasts. Intriguingly, we identified that many genes involved in immune responses, particularly those in specific C-C chemokine signaling pathways, were repressed with IMAP treatment. We next tested C-C chemokine ligands Ccl3, Ccl6, and Ccl17 in cardiac reprogramming and observed inhibitory effect on iCM formation, while corresponding inhibitors of Ccr1, Ccr4, and Ccr5 had the opposite effect. These results indicated that suppression of specific C-C chemokine signaling pathways was a direct down-stream event of IMAP treatment that enhanced cardiac reprogramming.

In conclusion, we identified a combination of four chemicals IMAP in suppressing specific C-C chemokine signaling pathways and facilitating MGT-induced cardiac reprogramming. Our studies revealed the role of these specific C-C chemokine signaling pathways in cardiac reprogramming and provide potential targets in iCM formation and its clinical applications.

---

Cardiovascular disease (CVD) has remained the leading cause of death in the United States and the whole world in the last decades(1). Myocardial infarction (MI) is the most severe result caused by CVD, which leads to impaired heart function because of massive cardiomyocyte (CM) loss and fibrosis(2). Existing treatments for MI are primarily pharmacological and device-based, and do not address the underlying problem of CM loss(3). Indeed, the prevalence of chronic cardiomyopathy is steadily increasing worldwide, escalating the urgency of developing novel therapies for this morbid disease.

Direct reprogramming of fibroblasts into CM-like cells by introducing three cardiac transcription factors Gata4, Mef2C and Tbx5 (G, M and T) has emerged as an attractive strategy to generate induced CMs (iCMs) (4-6). One major advantage of this strategy is utilizing abundant cardiac fibroblasts (CFs) in the heart to generate functional CMs to replace the scar tissue. Other advantages include the immunologic compatibility of the reprogrammed cells and lower possibility of tumorigenicity. However, low conversion rate, poor purity, and lack of precise conversion of iCMs still present significant challenges.

Several strategies have been applied to address these challenges in both mouse and human cells (5,7-27). It has been shown that adding other core transcriptional factors or protein kinases such as Hand2, Akt1 could further improve cardiac reprogramming efficiency in several fibroblast linages(5,12). Growth factors, such as fibroblast growth factor (FGF) 2, FGF10, and vascular endothelial growth factor (VEGF), can further promote cardiac reprogramming(17). Specific chemical compounds that influence different signaling pathways also enhance the efficiency and quality of cardiac reprogramming(16,26). Moreover, recent studies have identified epigenetic barriers in the reprogramming process. For example, targeting the polycomb ring finger oncogene Bmi1 and Mll1 H3K4 methyltransferase can enhance cardiac conversion(28,29). Single cell study of cardiac reprogramming indicates that RNA splicing also plays an important role in the cardiac reprogramming(27,30). Nevertheless, to bring this strategy closer to clinical studies, the iCM conversion and maturation need to be substantially improved.

Here we screened and identified four different chemical compounds, insulin growth factor-1 (IGF-1), Mll1 inhibitor MM589, transforming growth factor (TGF)-βinhibitor A83-01, and Bmi1 inhibitor PTC-209 (IMAP) that remarkably enhanced the conversion from fibroblasts into cardiomyocytes. We observed significant improvement in cardiac reprogramming including cardiac gene expression, sarcomere formation, calcium flux, and spontaneous beating. Importantly, RNA sequencing data showed that immune response related genes, especially those involved in chemokine signaling pathways, were downregulated. Therefore, we chose three chemokine axes, Ccl6-Ccr1, Ccl17-Ccr4, and Ccl3-Ccr5, as the potential targets. We then examined chemokine signaling pathway ligands Ccl3, Ccl6 and Ccl17 and observed decreased reprogramming efficiency. In contrast, corresponding chemokine receptor inhibitors significantly increased iCM formation, similar to IMAP treatment. Taken together, these findings identified specific immune responses as potential barriers for cardiac reprogramming and offered a promising strategy to resolve the low efficiency challenge of iCM formation by providing a novel chemical combination that targets immune responses.

## Results

### IGF-1, MM589, A83-01 and PTC-209 enhanced cardiac reprogramming

To identify small molecules that could enhance cardiac reprogramming, we used a screening platform reported previously(26,29). In this screening strategy, we isolated mouse embryonic fibroblasts (MEF) from a transgenic α-muscle heavy chain (α-MHC)-green fluorescent protein (GFP) reporter mouse(4). MEFs were treated with chemicals every three days after the infection of retrovirus expressing polycistronic Mef2c/Gata4/Tbx5 (MGT)(18). Two weeks later, the reprogramming efficiency was evaluated by percentage of GFP positive cells and related gene expression (Fig. 1A). Four chemical compounds, namely, insulin-like growth factor-1 (IGF-1), Mll1 inhibitor MM589, transforming growth factor (TGF)-β inhibitor A83-01, and Bmi1 inhibitor PTC-209, were identified that could improve cardiac reprogramming efficiency individually by 2 to 4 folds (Fig. 1B, C). Quantitative polymerase chain reaction (qPCR) indicated that each chemical played an important role in reprogramming by upregulation of cardiac related genes and/or downregulation of fibroblast related genes (Fig. 1D).

**Figure 1.**
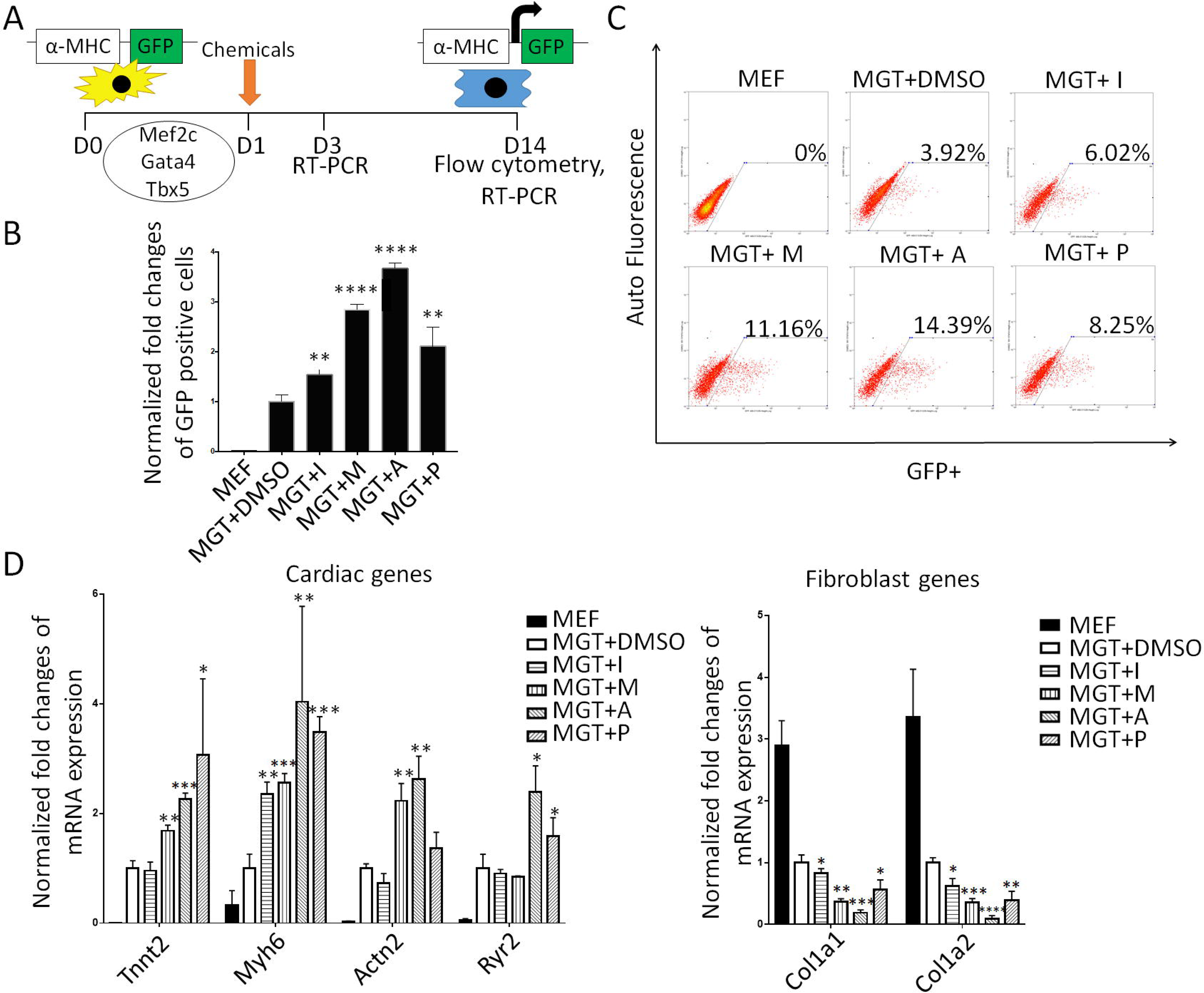
Drug screening identified four chemicals enhancing cardiac reprogramming. (A) A schematic diagram of the chemical screening using α-muscle heavy chain (α-MHC)-green fluorescent protein (GFP) mouse embryo fibroblasts (MEFs). (B) Bar graph showing % α-MHC-GFP+ cells two weeks after GMT infection with or without indicated chemical treatment. (C) Representative FACS plots showing iCMs with α-MHC-GFP+ cells after two weeks of reprogramming and quantification. (D) Bar graph representing major cardiac and fibroblast genes as determined by qPCR. Error bars indicate mean±s.e.m.; *P<0.05; **P<0.01; ***P<0.001; ****P<0.0001 compared with MGT+DMSO group.

### The combination of four chemicals IMAP achieved high reprogramming efficiency of MEFs

To explore the significance of each chemical in combination, we applied minus-one strategy to eliminate possible overlapping effects of these compounds. Using the same screening platform, we found that the combination of all four chemicals showed the highest efficiency and increased 6 folds of iCM formation compared to MGT+DMSO group (Fig. 2A). Eliminating every single compound in the IMAP caused a significant reduction of the reprogramming efficiency (Fig. 2B). qPCR results indicated that IMAP maximally enhanced the expression of cardiac genes and repressed fibroblast genes (Fig. 2C). After two weeks of reprogramming with IMAP treatment, immunofluorescence staining showed 3-4 folds increases in the expression of cardiac troponin T (cTnT) and α-actinin as well as significantly increased cTnT/GFP double positive cells (Fig. 2D, Supplementary Fig. S1A). Cell counting in several HPFs showed similar increase in cTnT, α-actinin, and cTnT/GFP double positive cells (Fig. 2E, Supplementary Fig. S1B).

**Figure 2.**
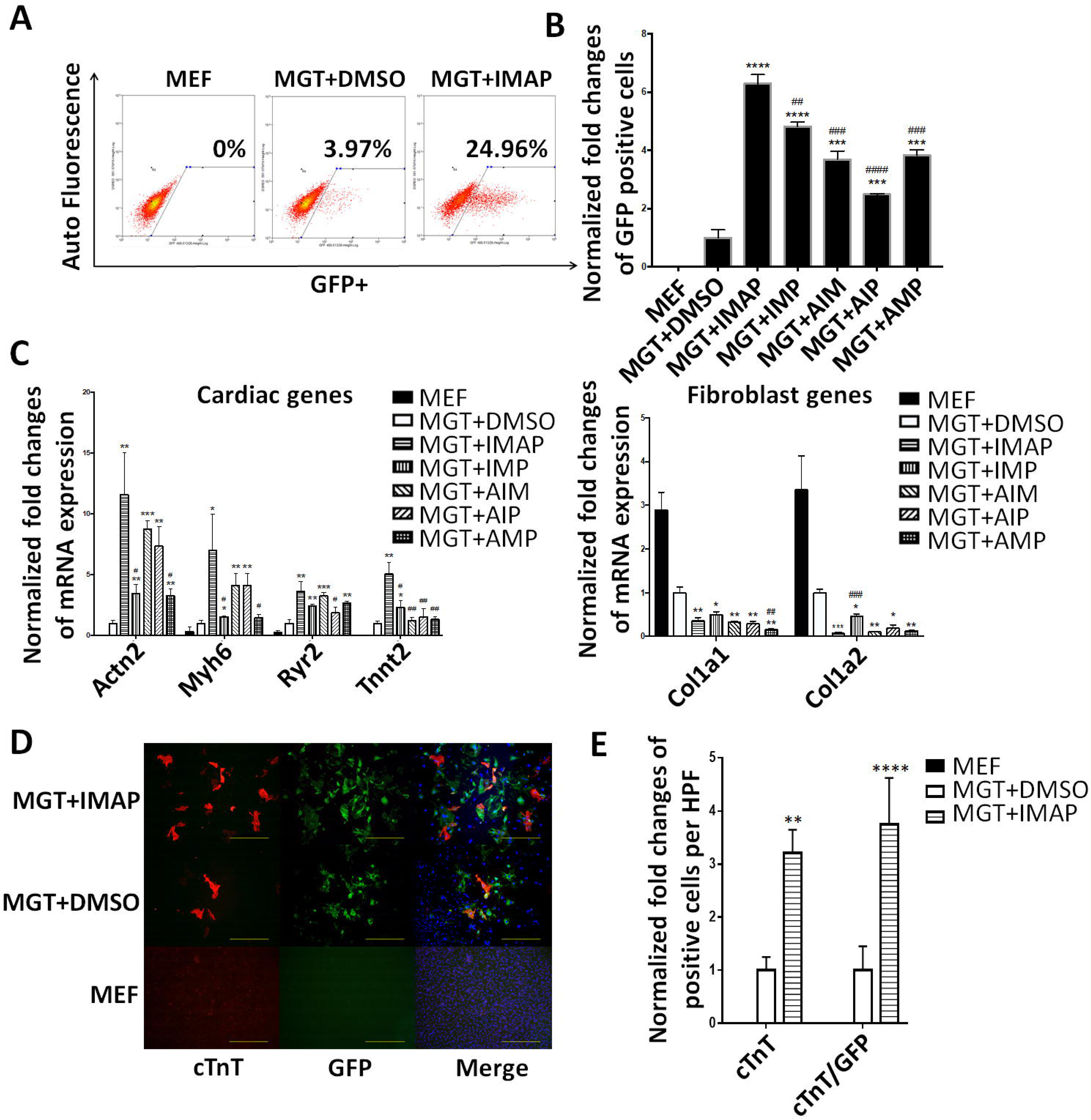
IMAP enhanced cardiac reprogramming of MEFs. (A) Representative FACS plots showing MEFs (Control) and iCMs with α-MHC-GFP+ cells after two weeks of reprogramming with IMAP or DMSO treatment. (B) Bar graph showing normalized fold changes of α-MHC-GFP+ cells two weeks after GMT infection with indicated chemical treatment. (C) Bar graph representing major cardiac and fibroblast genes as determined by qPCR in cells after two weeks of reprogramming and indicated chemical treatment. (D) Representative images of immunofluorescence staining and (E) Quantification of cardiac markers α-MHC-GFP (Green), cTnT (Red) and double positive cells in fibroblasts (Control) or cells treated with either MGT+DMSO or MGT+IMAP. (scale bar 500μm) Error bars indicate mean±s.e.m.; #P<0.05; ##P<0.01; ###P<0.001; ####P<0.0001 compared with MGT+IMAP group; *P<0.05; **P<0.01; ***P<0.001; ****P<0.0001 compared with MGT+DMSO group.

### IMAP also enhanced cardiac reprogramming of neonatal cardiac fibroblasts (NCFs)

Considering cardiac fibroblasts are the major in vivo target cell type for iCM reprogramming, we next determined the effect of IMAP on conversion of NCFs to iCMs. NCFs were isolated from the same α-MHC-GFP mouse line from which we obtained MEFs(31). Flow cytometry results indicated that the conversion efficiency was elevated 4-5 folds by IMAP compared with MGT+DMSO group (Fig. 3A). qPCR showed significant increase of cardiac genes and decrease of fibroblast related genes (Fig.3B). Immunofluorescence staining revealed a more significant increase of cTnT, α-actinin expression and cTnT/GFP double positive cells in NCFs compared with MEFs (Fig. 3C, D, Supplementary Fig. S1C-E) and clearly visualized sarcomere formation in iCMs derived from NCFs (Supplementary Fig. S1E).

**Figure 3.**
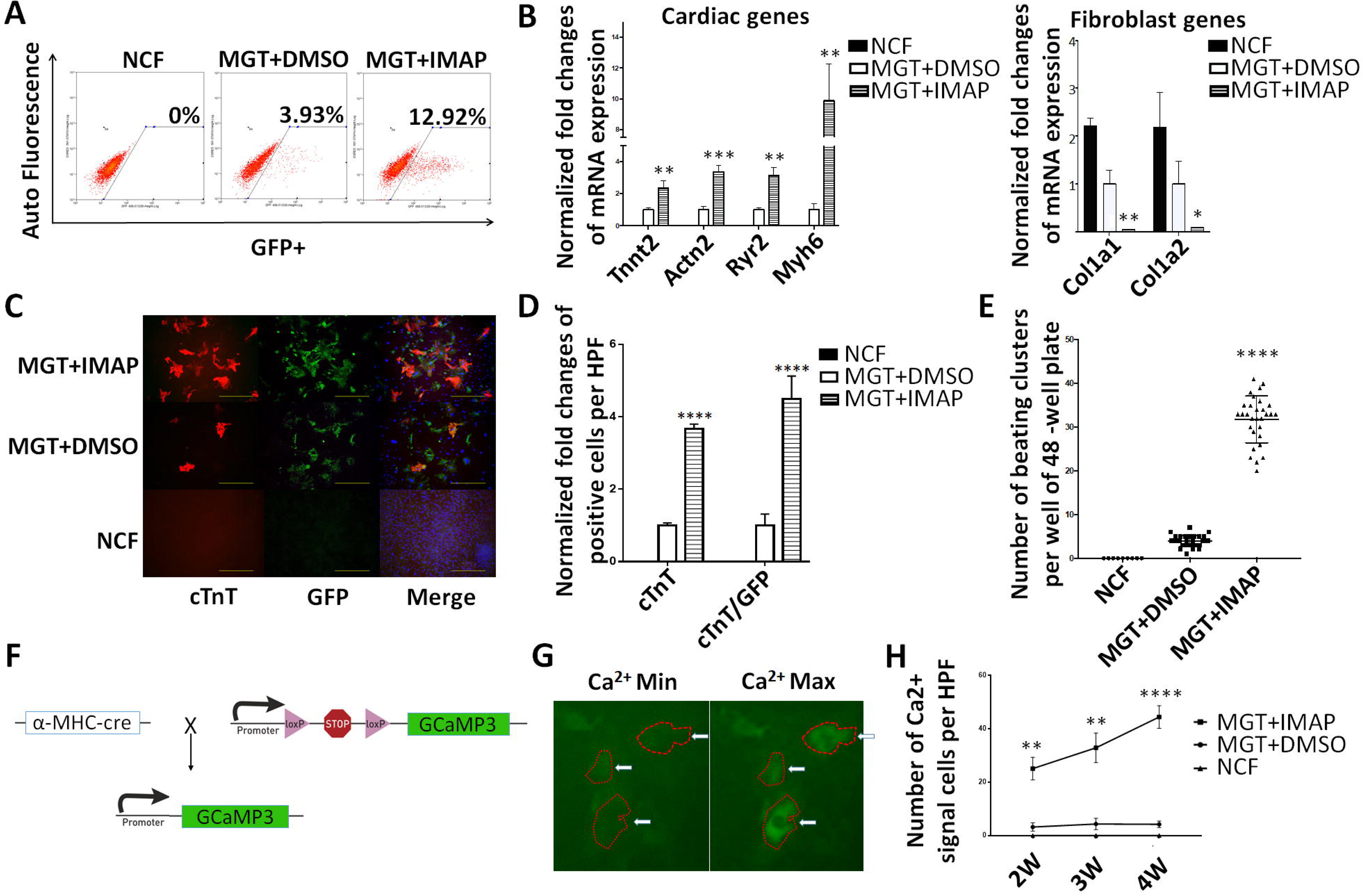
IMAP improved cardiac reprogramming efficiency of NCFs. (A) Representative FACS plots showing NCFs (Control) and iCMs with α-MHC-GFP+ cells after two weeks of reprogramming with MGT plus IMAP or DMSO treatment. (B) Bar graph representing major cardiac and fibroblast genes as determined by qPCR in cells after two weeks of reprogramming and indicated chemical treatment. (C) Representative images of immunofluorescence staining and (D)Quantification of cardiac markers α-MHC-GFP (Green), cTnT (Red) and double positive cells in NCFs (Control) or cells treated with either MGT+DMSO or MGT+IMAP.(scale bar 500μm) (E) Quantification of spontaneous beating iCMs per well of 48-well plate after two weeks of reprogramming and another two weeks with mature medium (n=10) (F) Schematic representation of the strategy for generating a calcium transient detection mouse line by crossing α-MHC-Cre mouse line with Rosa26A-Flox-Stop-Flox-GCaMP3 mouse line. (G) Representative images of iCMs exhibiting calcium transient. (H) Quantification of NCFs (Control) or iCMs with calcium transient activities after two weeks of reprogramming followed by another two weeks of mature medium treatment. (n=30 from 10 wells) Error bars indicate mean±s.e.m.; *P<0.05; **P<0.01; ***P<0.001; ****P<0.0001 compared with MGT+DMSO group.

Next, we tested spontaneous beating and calcium transient of iCMs since these two features represent characteristic cell functions of CMs. After treating NCFs with IMAP for two weeks and maturation medium for another two weeks, 8-9 fold increase in iCMs spontaneous beating was observed (Fig. 3E, Movie 1, 2). Additionally, spontaneous calcium transient was measured either by Ca2+ imaging with Rhod-3 staining (4) or calcium flux with GCaMP3 (7). With Rhod-3 staining in HPFs, hundreds of cells with spontaneous calcium transients were observed in MGT+IMAP group whereas only dozens were observed in MGT+DMSO group (Supplementary Fig. S1F, Movie 3, 4). To further measure calcium flux, αMHC-Cre/Rosa26A-Flox-Stop-Flox-GCaMP3 NCFs were used (Fig. 3F, G). IMAP promoted calcium flux as early as two weeks after treatment (Fig. 3H). Four-week treatment of IMAP resulted in 10-fold increase of calcium flux (Fig. 3H, Movie 5, 6), indicating that iCM can mature further under extended IMAP treatment.

Moreover, to determine if potential CM contamination could contribute to the iCM population, MEFs or NCFs without GMT infection and IMAP treatment were examined for GFP positive signals from initial fibroblast isolation to 4 weeks after iCM induction. With microscopic examination, flow cytometry or immunofluorescence staining (Fig. 2A-B, 2D-E, 3A, 3C-D, Supplementary Fig. S1C-E), no GFP signals were detected in those MEF or NCF control groups, indicating that CM contamination is highly unlikely in our experiments. Likewise, no cells in MEF or NCF control groups were observed to exhibit any spontaneous calcium transient or other CM properties (Fig. 3E, H, Supplementary Fig. S1F). All these results indicated that iCMs were reprogrammed from originally isolated fibroblasts and showed much better CM functionality in MGT+IMAP groups than those obtained from MGT+DMSO groups.

### RNA-sequencing data showed significant changes in gene expression profiles defining cell identity and immune responses

To identify systematically the downstream targets of IMAP in iCM formation, we first extracted the total RNAs and performed RNA-sequencing analyses using cells reprogrammed from α-MHC-GFP NCFs with two weeks of GMT and IMAP treatment. Cells reprogrammed with IMAP showed a much more robust cardiomyocyte gene expression profiles compared with MGT+DMSO group (Fig. 4A). iCMs derived from MEFs showed similar results with 1442 differentially expressed genes when comparing MGT+IMAP group with MGT+DMSO group (Fig. 4B, Supplementary Fig. S2A). 528 overlapping genes were identified in both datasets, and gene ontology (GO) enrichment analysis revealed that IMAP stimulated gene expression related to muscle structure formation and muscle contraction as well as decreased gene expression in extracellular matrix formation (Fig. 4B, Supplementary Fig. S2B, C). Intriguingly, GO analysis identified significant gene repression in immune system process and inflammation response by IMAP (Fig. 4B, Supplementary Fig. S2C). We next examined in more detail the effect of IMAP on immune response gene expression. Several immune response representative genes were tested in reprogramming. Il6, Tnf, Ccl2 and Ptgs2 expression decreased as early as day 3 after IMAP treatment (Fig. 4C). RNA-seq analysis further identified many immune related genes that were significantly downregulated after IMAP treatment in both NCFs (Fig. 4D) and MEFs (Supplementary Fig. S2D). GO molecular function analysis revealed that these downregulated genes shared similar biological function in C-C chemokine signaling pathways (Fig. 4E, Supplementary Fig. S2E). These data suggested that C-C chemokine signaling pathways could be potential novel targets in improving iCM formation and may provide mechanistic insights into IMAP-mediated cardiac reprogramming.

**Figure 4.**
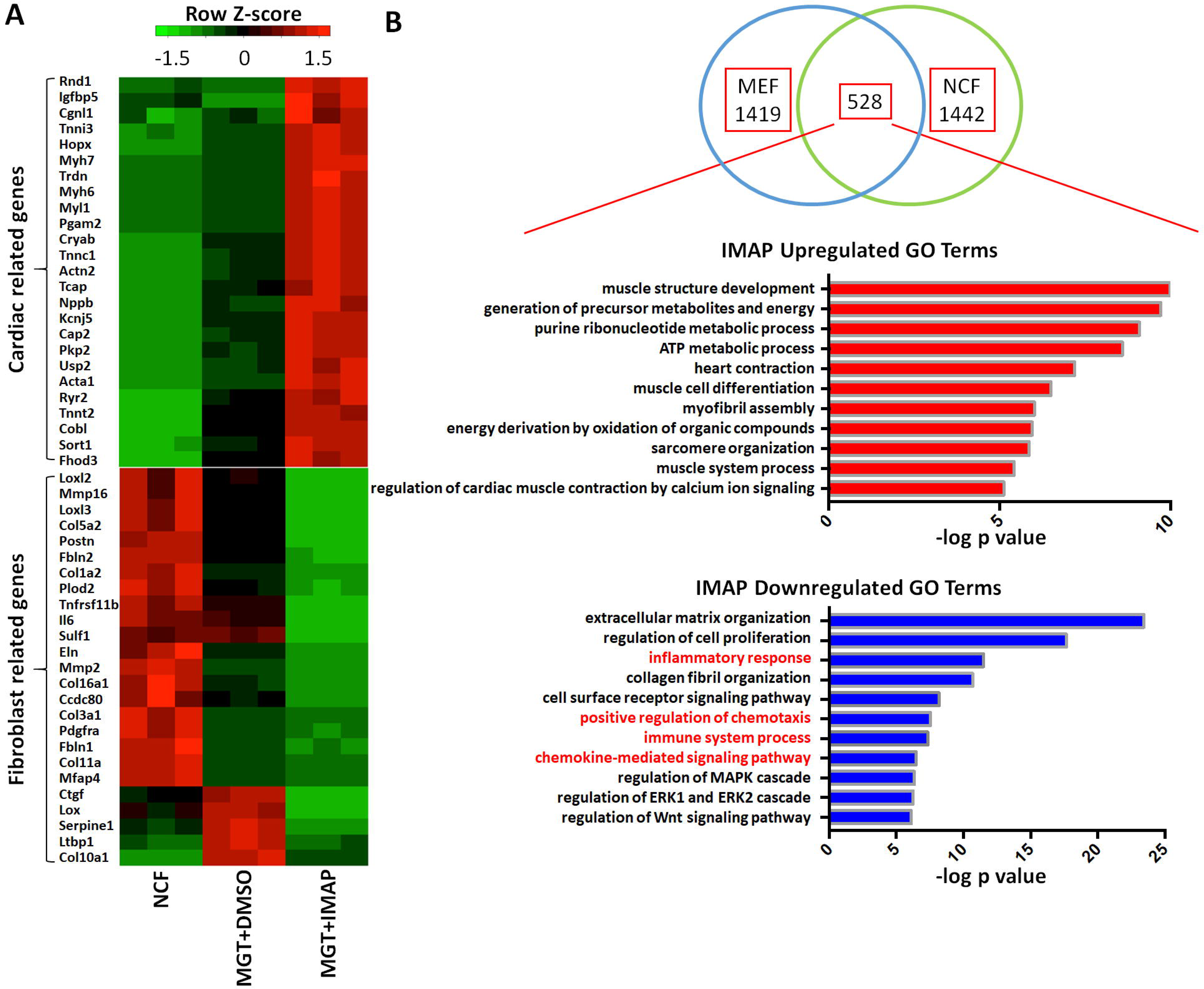
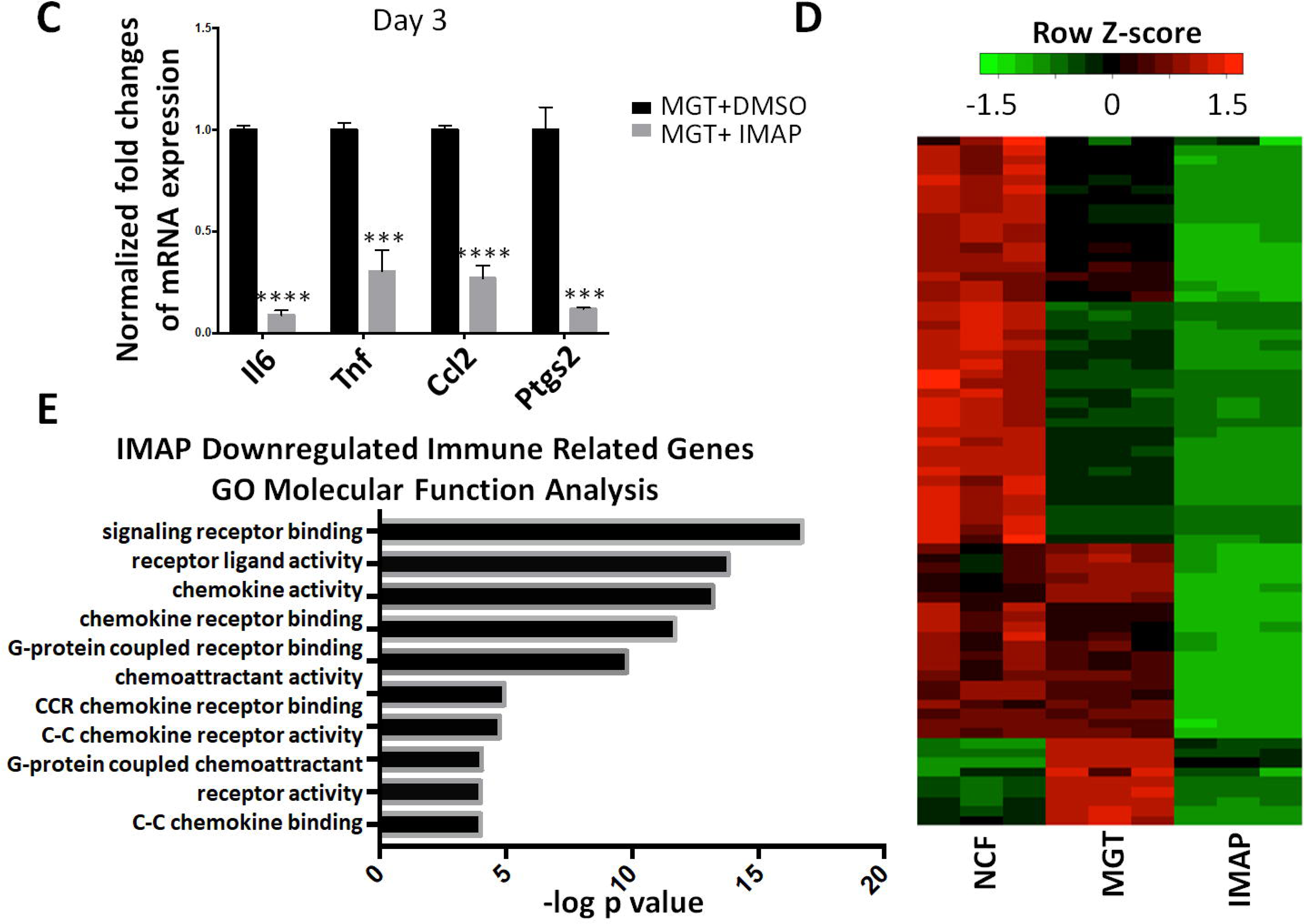
RNA-sequencing data showed significant changes of gene expression profiles defining cell identity and immune responses. (A) Heat map showing differential expression of representative cardiac and fibroblast related genes among NCFs (Control), MGT+DMSO and MGT+IMAP group. (B) Venn diagram showing the number of differentially expressed genes between MGT+DMSO and MGT+IMAP group in MEFs and NCFs. Bar graph showing the top gene ontology (GO) terms of the upregulated (red) or downregulated (blue) genes expressed between MGT+DMSO group and MGT+IMAP group. (C) Bar graph showing representative immune response related gene expression between MGT+DMSO group and MGT+IMAP group at day 3 after MGT infection. (D) Heat map showing differential expression of immune related genes among NCFs (Control), MGT+DMSO and MGT+IMAP group. (E) Bar graph for gene ontology (GO) molecular function analysis terms of IMAP down-regulated immune related genes compared with MGT+DMSO group. Error bars indicate mean±s.e.m.; ***P<0.001; ****P<0.0001 compared with MGT+DMSO group.

### Specific C-C Chemokine Signaling Pathways were likely important barriers in cardiac reprogramming that were overcome by IMAP treatment

Based on the above results, we examined C-C chemokine signaling pathways Ccl6-Ccr1, Ccl17-Ccr4 and Ccl3-Ccr5 in cardiac reprogramming as they were significantly downregulated in MGT+IMAP treated group. We first performed loss-of-function studies by testing the inhibitors of corresponding C-C chemokine receptors in cardiac reprogramming. We applied Ccr1 inhibitor BX471, a CCR4 antagonist, and Ccr5 inhibitor Maraviroc in α-MHC-GFP MEFs. In MGT-induced cardiac reprogramming, all three inhibitors enhanced reprogramming individually while the three inhibitors together, called 3i, achieved a combinatorial improvement (Fig. 5A). These chemokine receptor inhibitors promoted the expression of cardiac-related genes and repressed the expression of fibroblast-related genes, similar to the effect of IMAP (Fig. 5B). Similar results were observed from NCF reprogramming (Fig. 5C, D). Immunofluorescence staining showed much higher number of cTnT, α-actinin and cTnT/GFP double positive cells in MGT+3i treated group compared with MGT+DMSO group (Fig. 5E, F, Supplementary Fig. S3A, B).

**Figure 5.**
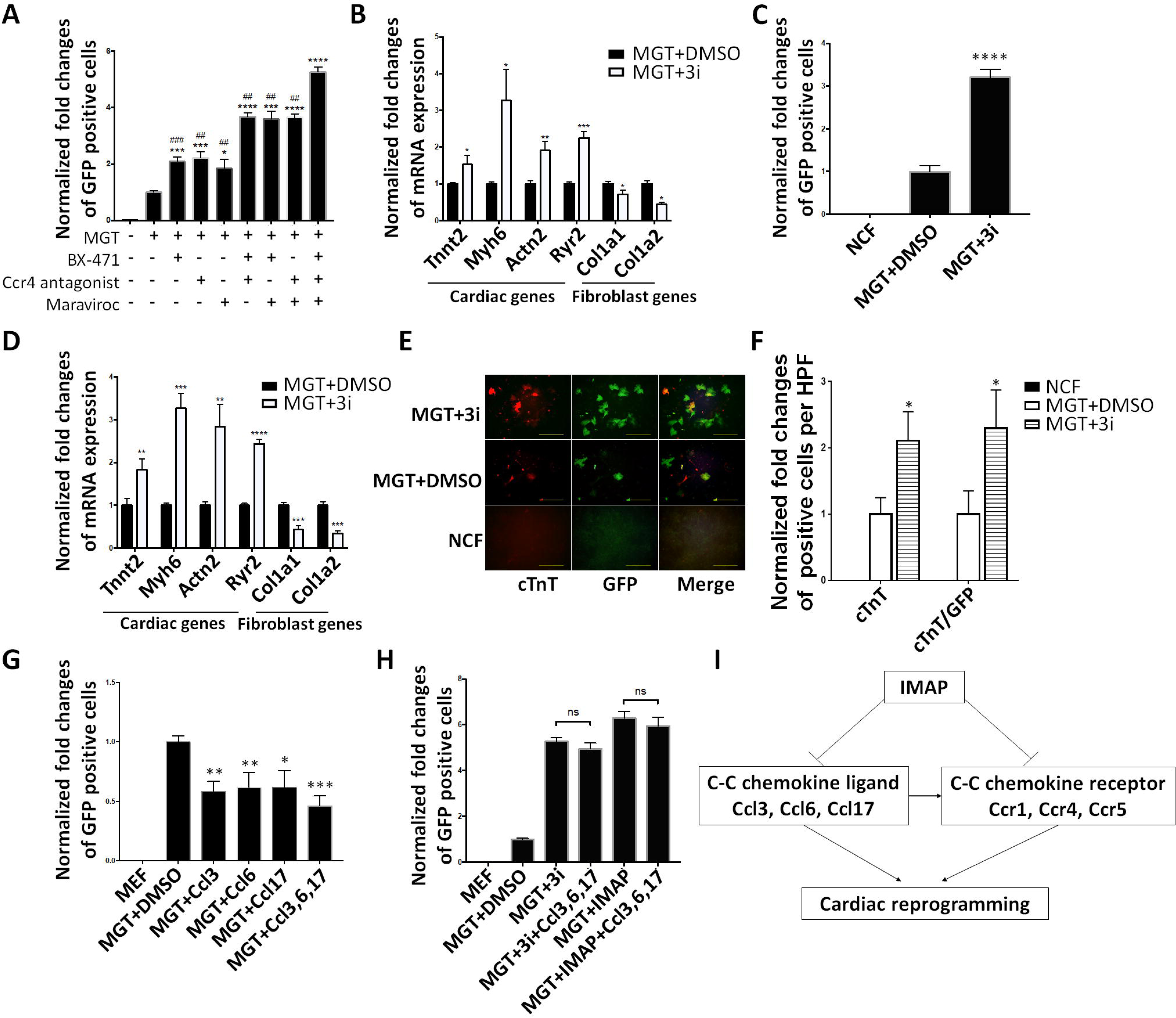
Specific C-C Chemokine Signaling Pathways were likely important barriers in cardiac reprogramming that were overcome by IMAP treatment. (A) Bar graph showing normalized fold changes of α-MHC-GFP+ cells two weeks after GMT infection with indicated chemical treatment in MEFs and (C) NCFs. (B) Bar graph representing major cardiac and fibroblast genes as determined by qPCR in cells after two weeks of reprogramming and indicated chemical treatment in MEFs and (D) NCFs. (E) Representative images of immunofluorescence staining and (F) quantification of cardiac markers α-MHC-GFP (Green), cTnT (Red) and double positive cells in NCFs (Control) or cells treated with either MGT+DMSO or MGT+3i. (G) Bar graph showing normalized fold changes of α-MHC-GFP+ cells two weeks after GMT infection with indicated C-C chemokine ligands treatment in MEFs. (scale bar 500μm) (H) Bar graph showing normalized fold changes of α-MHC-GFP+ cells two weeks after GMT infection with indicated with indicated treatment in MEFs. (I) A schematic diagram showing IMAP-enhanced cardiac reprogramming by suppressing the expression of several specific C-C chemokine ligands and receptors. Error bars indicate mean±s.e.m.; #P<0.05; ##P<0.01; ###P<0.001; ####P<0.0001 compared with MGT+3i group; *P<0.05; **P<0.01; ***P<0.001; ****P<0.0001 compared with MGT+DMSO group.

To further validate our findings that blocking the activities of specific C-C chemokine signaling pathways increased iCM reprogramming, we also performed gain-of-function experiments to examine the effect of chemokine ligands in reprogramming. Addition of chemokine ligands Ccl3, 6, and 17 in MGT-mediated reprogramming significantly reduced reprogramming efficiency (Fig. 5G, Supplementary Fig. S3C). In contrast, the inhibitory effect of these ligands was abolished when their receptor expressions or activities were inhibited in the presence of IMAP and 3i (Fig. 5H, Supplementary Fig. S3D), indicating that these ligands’ inhibitory effect on reprogramming was specifically through C-C chemokine signaling pathways. Taken together, our research showed that IMAP enhanced cardiac reprogramming by eliminating iCM reprogramming barriers imposed by specific C-C chemokine pathways (Fig. 5I).

### Discussion

We report here a chemical cocktail that enhances cardiac reprogramming by suppressing specific C-C chemokine signaling pathways. Administration of chemicals IGF-1, MM589, A83-01 and PTC-209, named IMAP, significantly enhances the reprogramming efficiency of MEFs or NCFs to iCMs. IMAP promotes cardiac gene expression, sarcomere formation, spontaneous beating and calcium transient in reprogramming, indicating a significant role of IMAP in promoting maturation and functionality of iCMs. Importantly, repression of Ccr1, Ccr4 and Ccr5 signaling pathways is likely to be responsible for cardiac reprograming augmentation mediated by IMAP. Direct inhibition of these chemokine receptors has achieved similar positive effect on cardiac reprogramming whereas their corresponding ligands repress reprogramming. Therefore, our study provides a novel platform of identifying chemicals that repress Specific C-C chemokine signaling pathways for cardiac reprogramming.

In this study, we have developed a new platform enhancing cardiac reprogramming by combining four chemicals named IMAP. Two chemicals IGF-1(12) and A83-01 (13), targeting IGF growth factor and TGF-β signaling, enhance reprogramming, as reported previously. PTC-209 is an inhibitor of polycomb ring finger oncogene Bmi1 which functions through chromatin remodeling as an essential epigenetic regulator(32) and has been identified as a barrier in cardiac reprogramming(28). Importantly, our previous work reveals that targeting WD repeat domain 5 (WDR5) and blocking the WDR5–mixed lineage leukemia (MLL) protein–protein interaction enhance cardiac reprogramming(29). In this study, we have identified that MM589, a newly invented version of WDR5-MLL1 inhibitor, promotes reprogramming. IMAP combination shows significant enhancement of MGT-induced cardiac reprogramming and elimination of each of them decreases reprogramming efficiency. Our data thus indicate that these four chemicals work together to convert fibroblasts into iCMs.

Our IMAP studies identify an unexpected connection between C-C chemokine signaling pathways and cardiac reprogramming. In particular, we have observed that immune response related genes, especially those involved in chemokine signaling pathways, were downregulated in IMAP-mediated cardiac reprogramming. Subsequently, we have identified three chemokine axes, Ccl6-Ccr1, Ccl17-Ccr4, and Ccl3-Ccr5, as important reprogramming barriers. Administration of these three chemokine signaling pathway ligands Ccl3, Ccl6, and Ccl17 significantly decreases reprogramming efficiency. In contrast, addition of corresponding chemokine receptor inhibitors in our reprogramming assay significantly increased iCM formation, similar to IMAP treatment. Thus, our study reveals that immune responses may act as vital barriers for cardiac reprogramming. Mechanistically, the effect of chemokine signaling pathways in reprogramming could be linked to pro-fibrotic signaling, which has been identified to inhibit cardiac reprogramming as shown by others (13,15) and our studies. The three C-C chemokine pathways we have identified are all reported to promote fibrosis process both in rodents and human (33-35). In addition, the effect of C-C chemokine pathways could be through Wnt pathway regulation. Wnt pathway-related genes also show significant decrease in our MGT+IMAP treatment group compared with MGT+DMSO group, and Wnt pathway is required for chemokine signaling-mediated functions in cardiac reprogramming (26,36). Future studies will be required to further reveal the underlying mechanisms.

Even though our identification of chemokine signaling pathways in inhibiting cardiac reprogramming is surprising, studies of immune response and chemokine signaling pathways in cell fate change and development provide strong rationale that chemokine signaling pathways are indeed very likely a key barrier in cardiac reprogramming. Traditionally, C-C chemokine signaling pathways were initially considered merely as low-molecular-weight proteins that stimulate recruitment of leukocytes(37). However, recent studies show that specific C-C chemokine signaling pathways, such as stromal cell-derived factor-1-cxcr4 axis, can exerts negative feedback on the Fgf pathway at the beginning of fin regeneration(38) and regulate myoD and myf5 expression and fast fiber formation in muscle differentiation of zebrafish(39). Importantly, C-C chemokine signaling pathways can significantly impact cardiac development. In particular, Cxcr4 is considered as an important marker for cardiomyocyte progenitors and may play a functional role in their differentiation(40). Further studies also show that repressing Cxcl10 and Ccl5 by WT1 is also required during cardiac development while Cxcr4 is required for spontaneous beating of hiPSC-CMs (41,42). In addition, targeting C-C chemokine signaling pathways has been proposed in cardiac repair after injury (43). Finally, anti-inflammatory drugs also facilitate induced pluripotent stem cell reprogramming(44,45), and anti-inflammation treatment may benefit cardiac reprogramming (25). Therefore, further investigation of chemokine pathways in cardiac reprogramming may provide key insights into cell fate change.

Direct cardiac reprogramming provides a promising strategy for repairing injured heart after massive cardiomyocyte loss. Application of chemical approaches for reprogramming represents a highly desirable approach to improve the efficiency of iCM formation, as reported previously by us (29) and others (13,15,16,21,26,46,47). Importantly, our studies have identified specific C-C chemokine signaling pathways regulated by a combination of four chemicals IMAP in cardiac reprogramming. We expect that future studies towards combined chemical approaches in targeting chemokine signaling pathways will further improve the iCM formation and bring cardiac reprogramming closer to clinical applications.

## Materials and Methods

### Mouse lines

The α-MHC-GFP transgenic mice were used to derive MEFs and NCFs as described previously(31,48). All animal related procedures were approved by the Institutional Animal Care and Use Committee of the University of Michigan and are consistent with the National Institutes of Health Guide for Use and Care of Animals.

### Plasmids

The pMXs based retroviral polycistronic vector encoding Mef2c, Gata4, Tbx5 were provided from Dr. Li Qian’s lab^(18)^. This polycistronic constructs vector was constructed by DNA fragment containing Mef2c, Gata4 and Tbx5 sequentially which were separated by oligonucleotides encoding P2A and T2A peptides. This polycistronic vector also contains puromycin selection marker for cell purification.

### Primary cell isolation

Preparation of MEFs (isolated at E13.5) was previously reported(48). Briefly, embryos were harvested from α-MHC-GFP transgenic mice at 13.5 days post coitum followed by decapitation and removal of internal organs. The tissue was minced and digested with 0.05% trypsin/EDTA (Gibco, Thermo Fisher Scientific). Cells were resuspended in MEFs medium (DMEM medium containing 10%FBS, 1% penicillin/ streptomycin, 10 μl/ml GlutaMAX and 1 mM Sodium Pyruvate) and plated onto one 10cm dish per embryo. Cells were passaged at the ratio of 1:3 (passage 1). Passage 3 MEFs were used for reprogramming.

NCFs were isolated from P2-P3 α-MHC-GFP transgenic or αMHC-Cre/Rosa26A-Flox-Stop-Flox-GCaMP3 mice as described previously(31). Briefly, heart tissue was isolated, minced and digested with 0.05% Trypsin-EDTA. Then NCFs were collected with type II collagenase (0.5 mg/ml) in HBSS. After washing and resuspending in DPBS with 2.5 g BSA and 0.5M EDTA, cells were incubated with CD90.2 microBeads (miltenyibiotec, Midi MACS Starting Kit) at 4°C for 30min. Positive cells were isolated by Magnetic-activated cell sorting (MACS) and plated onto a 10cm dish with FB medium (IMDM media with 20%FBS and 1% penicillin/ streptomycin) for future use. Isolated fibroblasts were routinely examined under fluorescence microscope or with GFP immunostaining to determine potential CM contamination.

### Chemicals information

IGF-1was from Peprotech (100-11), MM589 was from Dr. Shaomeng Wang’s lab, A83-01 was from Stemgent (04-0014), PTC-209 was from Sigma (SML1143-5MG). C-C chemokine receptor inhibitors including Ccr1 inhibitor BX471, CCR4 Antagonist, and Ccr5 inhibitor Maraviroc were from Cayman (18503, 21885,14641). Recombinant Mouse Ccl3, Ccl6, and Ccl17 were from Biolegend (593802, 758502, 581402).

### Retrovirus preparation

80% confluent 10 cm plates of Plat-E cells were transfected by using Lipofectamine 2000 (Thermo Fisher Scientific) with 10 μg retrovirus vectors in additional 1.5 ml Opti-MEM (Thermo Fisher Scientific). 24h later, medium was changed into 10ml fresh MEFs medium. Forty-eight hours and seventy-two hours after transfection, viral medium was collected twice and filtered through a 0.45-mmcellulose filter. The virus containing medium was added 1/5 vol of 40% PEG8000 solution to make a final concentration of 8% PEG8000. The mixture was kept at 4 °C overnight and spun at 3000 g, 4 °C, 30 min to get concentrated. Virus was resuspended by fresh MEFs medium with 8 μg/ml polybrene (Sigma).

### Direct reprogramming of fibroblasts to iCMs

The protocol of direct cardiac reprogramming was similar to previous studies with minor optimization(31). Fresh fibroblasts were seeded on tissue culture dishes at a density of 10,000 cells/cm^2^ before virus infection. Fibroblasts were infected with freshly made viral mixture containing 8 μg/ml polybrene (Sigma) 24 h post seeding. Twenty-four hours later, the viral medium was them replaced with induction medium composed of DMEM/199 (4:1) (Gibco, Thermo Fisher Scientific) containing 10% FBS, 1% penicillin/ streptomycin and 10 μl/ml GlutaMAX (Gibco, Thermo Fisher Scientific). Medium was changed every 2–3 days with or without indicated chemicals for two weeks before cells were examined. 1 μg/ml puromycin (SIGP8833-25MG, Sigma) was added into the medium 3 days after infection to eliminate none infected cells. For spontaneous beating and calcium transient assessment experiments, induction medium was replaced by mature medium containing StemPro-34 SF medium (Gibco, Thermo Fisher Scientific), GlutaMAX (10 μl/ml, Gibco, Thermo Fisher Scientific), ascorbic acid (50 μg/ml, Sigma), recombinant human VEGF165 (5 ng/ml, R&D Systems), recombinant human FGF basic146 aa (10 ng/ml, R&D Systems), and recombinant human FGF10 (25 ng/ml, R&D Systems). Then mature medium was changed every 2–3 days for another 2 weeks as previously described(17).

### Immunocytochemistry

Cells were fixed with 4% formaldehyde for 15min and permeabilized with 0.1% Triton X-100 in PBS for 15min at room temperature. Cells were blocked with 4% goat serum in PBS for 1 hr and then incubated with primary antibodies against cTnT (Thermo Fisher Scientific), α-actinin (Abcam), and GFP (Thermo Fisher Scientific) overnight at 4°Cfollowed by incubation with appropriate Alexa fluorogenic secondary antibodies (Thermo Fisher Scientific) at room temperature for 1 hr. cTnT, α-actinin, and cTnT/GFP double positive cells were manually quantified by single-blind method from ten randomly HPFs within each well.

### Flow cytometry

Fluorescence flow cytometry data was from 10000 single cell events which were collected using a standard MoFloAstrios flow cytometer (Immunocytometry Systems; Becton Dickinson, Detroit, MI, USA). Data was analyzed using Summit (Becton Dickinson).

### Quantitative real time PCR (qPCR)

Total RNAs from all induced cells with two-week GMT and IMAP treatment were extracted using Trizol Reagent (Thermo Fisher Scientific) following the manufacturer’s instructions. RNA integrity was determined using formaldehyde denaturalization agarose gel electrophoresis. RNA concentrations were measured with the Nanodrop spectrophotometer (Thermo Fisher Scientific). RNA was reverse transcribed using iScript cDNA Synthesis Kit (BioRad). qPCR was performed using StepOne

Real-Time PCR System (Thermo Fisher Scientific). Primer oligonucleotides were synthesized by Sigma and are listed in Supplementary Table S1.

### Spontaneous beating and calcium transient assessment

Spontaneous beating assessment was performed by light microscopy at room temperature after indicated treatment of NCFs at four-week time point. Beating cell number was manually quantified by single-blind method after isoproterenol treatment in each well of 48-well plate.

Calcium transient was measured with Rhod-3 Calcium Imaging Kit (Thermo Fisher Scientific) according to the manufacturer’s instructions. αMHC-Cre/Rosa26A-Flox-Stop-Flox-Gcamp3 NCFs were also used for calcium transient measurements as previously reported (12) which were performed by fluorescence microscopy at room temperature after the indicated treatment of retrovirus and chemicals. Three HPFs of view were randomly selected within each well, video recorded for 3 min and manually quantified.

### RNA-sequencing

Total RNAs from all induced cells with two-week GMT and IMAP treatment were isolated using TRIzol following the provider’s instructions. RNA (RIN > 8.5) was used for RNA-seq library preparation using NEBNext® Ultra™ II Directional RNA Library Prep Kit for Illumina (E7760S). The libraries were sequenced using Hiseq 4000 by the University of Michigan Sequencing Core. The quantification of RNA expression was estimated by Kallisto (49). Differential gene expression analysis was done using the R package DESeq2. The abundance of genes was used to calculate fold change and p values. Cutoff values of fold change greater than 2 and p values less than 0.05 were then used to select for differentially expressed genes between sample group comparisons.

Significant pathway enrichment analysis was performed using PANTHER Overrepresentation Test (release 20150430, http://geneontology.org).

### Statistic analysis

Results were presented as mean ±s.e.m. Statistical difference between groups was analyzed by one-way ANOVA followed by the Student– Newman–Keuls multiple comparisons tests. A P-value <0.05 was regarded as significant. Each experiment was performed at least twice.

## Acknowledgements

This work was supported by the National Institutes of Health Grant HL109054, a Pilot Award from the Joint Institute of University of Michigan Health System and Peking University Health Science Center (to WZ).

