## Supplemental Table S1, Figure S1-3 for "Enhancing Cardiac Reprogramming by Suppressing Specific C-C Chemokine Signaling Pathways"

**Supplement**

Table S1. Primers used in qPCR.

| Genes | Forward Primer | Backward Primer |
|---|---|---|
| Tnnt2 | CTGAGACAGAGGAGGCCAAC | ACCAAGTTGGGCATGAAGAG |
| Actn2 | CATCGAGGAGGATTTCAGGAAC | CAATCTTGTGGAACCGCATTTT |
| Myh6 | GCCCAGTACCTCCGAAAGTC | GCCTTAACATACTCCTCCTTGTC |
| Ryr2 | ACGGCGACCATCCACAAAG | AAAGTCTGTTGCCAAATCCTTCT |
| Col1a1 | GGTGAGCCTGGTCAAACGG | ACTGTGTCCTTTCACGCCTTT |
| Col1a2 | GCTCCTCTTAGGGGCCACT | CCACGTCTCACCATTGGGG |

Figure S1. IMAP-enhanced cardiac reprogramming in both MEFs and NCFs
(A) Representative images of immunofluorescence staining and (B) Quantification of cardiac markers α-MHC-GFP (Green) and α-actinin (Red) in fibroblasts (Control) or cells treated with either MGT+DMSO or MGT+IMAP for two weeks. (scale bar 500μm) (C) Representative images of immunofluorescence staining including enlarged image showing sarcomere formation (scale bar 200μm) of iCMs and (D)Quantification of cardiac markers α-MHC-GFP (Green) and α-actinin (Red) in NCFs (Control) or cells treated with either MGT+DMSO or MGT+IMAP for two weeks. (scale bar 500μm) (E) Representative images of immunofluorescence staining of cardiac markers α-actinin and cTnT treated with either MGT+DMSO or MGT+IMAP in NCFs. (scale bar 2.0mm) (F) Quantification of NCFs (Control) or iCMs with calcium transient activities stained by Rhod-3 dye after two weeks of reprogramming followed by another two weeks of mature medium treatment. Error bars indicate mean±s.e.m.; $^{**}P<0.01$, $^{****}P<0.0001$ compared with MGT+DMSO group.

Figure S2. RNA-sequencing data in MEFs showed significant changes of gene expression profiles
(A) Heat map showing differential expression of representative cardiac and fibroblast related genes among MEFs (Control), MGT+DMSO and MGT+IMAP group. (B) Bar graph showing the top gene ontology (GO) terms of the upregulated genes between MGT+DMSO group and MGT+IMAP group. (C) Bar graph showing the top GO terms of the downregulated genes between MGT+DMSO group and MGT+IMAP group. (D) Heat map showing representative immune response related genes expression between MGT+DMSO group and MGT+IMAP group two weeks after MGT infection. (E) Bar graph for GO molecular function analysis terms of IMAP down-regulated immune related genes compared with MGT+DMSO group.

Figure S3. IMAP enhance cardiac reprogramming by overcoming the barriers of specific C-C chemokine signaling pathways
(A) Representative images of immunofluorescence staining and (B)Quantification of cardiac markers α-MHC-GFP (Green) and α-actinin (Red) in NCFs (Control) or cells treated with either MGT+DMSO or MGT+3i for two weeks. (scale bar 500μm) (C) Bar graph representing major cardiac and fibroblast genes as determined by qPCR in MEFs after two weeks of reprogramming and indicated C-C chemokine ligands treatment. (D) Bar graph representing major cardiac and fibroblast genes as determined by qPCR in MEFs with indicated chemicals and C-C chemokine ligands treatment. Error bars indicate mean±s.e.m.; $^{*}P<0.05$; $^{**}P<0.01$; $^{***}P<0.001$ compared with MGT+DMSO group.

Movie 1. Showing spontaneous beating of iCMs derived from neonatal cardiac fibroblasts (NCFs) in MGT+IMAP group group;

Movie 2. Showing spontaneous beating of iCMs derived from NCFs in MGT+DMSO group group;

Movie 3. Showing calcium transient activities in MGT+IMAP group using Rhod3 staining;

Movie 4. Showing calcium transient activities in MGT+DMSO group using Rhod3 staining;

Movie 5. Showing calcium transient activities in MGT+IMAP group using αMHC-Cre/Rosa26A-Flox-Stop-Flox-GCaMP3 NCFs.

Movie 6. Showing calcium transient activities in MGT+DMSO group using αMHC-Cre/Rosa26A-Flox-Stop-Flox-GCaMP3 NCFs.

Figure S1

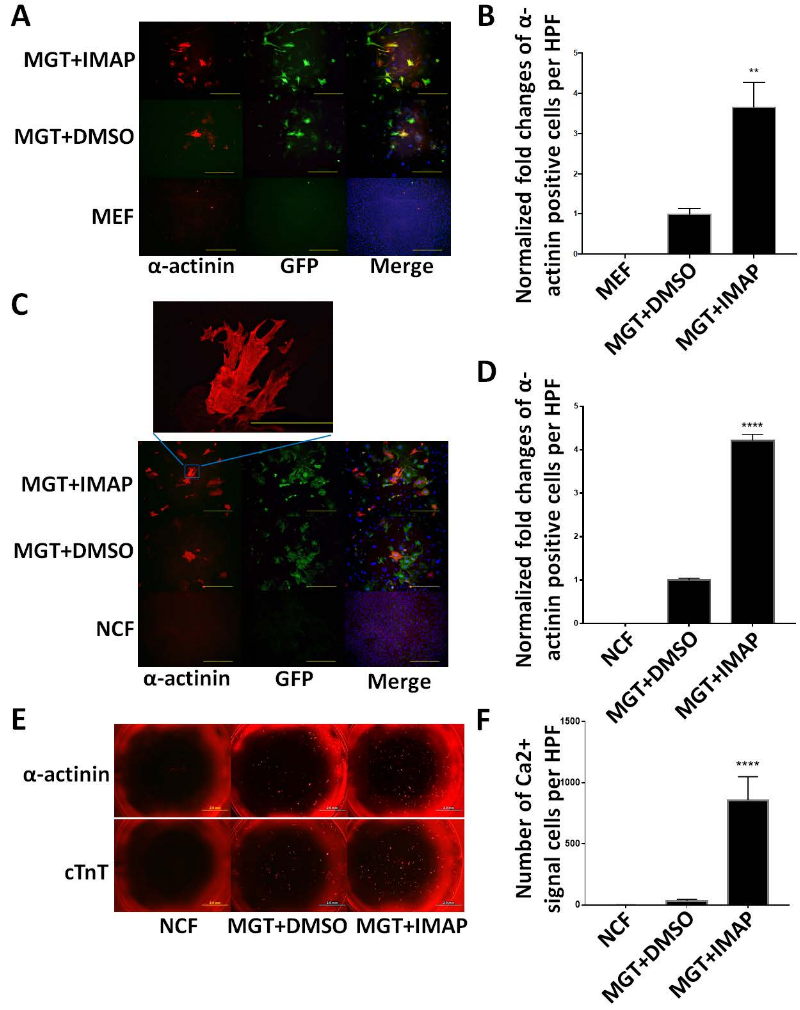
A
MGT+IMAP
MGT+DMSO
MEF
α-actinin
GFP
Merge
B
Normalized fold changes of α-actinin positive cells per HPF
5
4
3
2
1
0
**
MEF
MGT+DMSO
MGT+IMAP
C
MGT+IMAP
MGT+DMSO
NCF
α-actinin
GFP
Merge
D
Normalized fold changes of α-actinin positive cells per HPF
5
4
3
2
1
0
****
NCF
MGT+DMSO
MGT+IMAP
E
α-actinin
cTnT
NCF
MGT+DMSO
MGT+IMAP
F
Number of Ca2+ signal cells per HPF
1500
1000
500
0
****
NCF
MGT+DMSO
MGT+IMAP

Figure S2

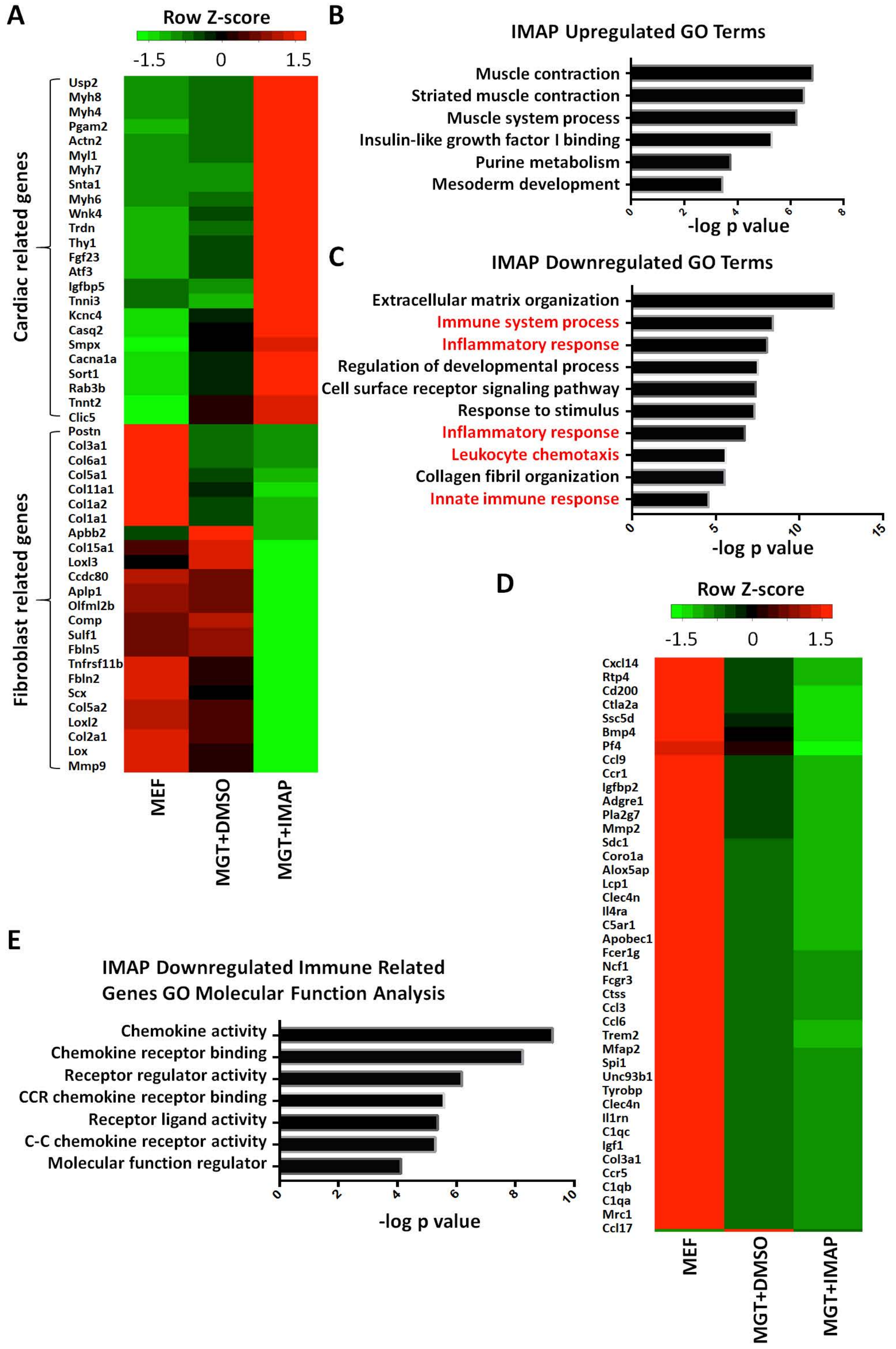
A
Row Z-score
-1.5
0
1.5
Cardiac related genes
Usp2
Myh8
Myh4
Pgam2
Actn2
Myl1
Myh7
Snta1
Myh6
Wnk4
Trdn
Thy1
Fgf23
Atf3
Igfbp5
Tnni3
Kcnc4
Casq2
Smpx
Cacna1a
Sort1
Rab3b
Tnnt2
Clic5
Fibroblast related genes
Postn
Col3a1
Col6a1
Col5a1
Col11a1
Col1a2
Col1a1
Apbb2
Col15a1
Loxl3
Ccdc80
Aplp1
Olfml2b
Comp
Sulf1
Fbln5
Tnfrsf11b
Fbln2
Scx
Col5a2
Loxl2
Col2a1
Lox
Mmp9
MEF
MGT+DMSO
MGT+IMAP
B
IMAP Upregulated GO Terms
Muscle contraction
Striated muscle contraction
Muscle system process
Insulin-like growth factor I binding
Purine metabolism
Mesoderm development
0
2
4
6
8
-log p value
C
IMAP Downregulated GO Terms
Extracellular matrix organization
Immune system process
Inflammatory response
Regulation of developmental process
Cell surface receptor signaling pathway
Response to stimulus
Inflammatory response
Leukocyte chemotaxis
Collagen fibril organization
Innate immune response
0
5
10
15
-log p value
D
Row Z-score
-1.5
0
1.5
Cxcl14
Rtp4
Cd200
Ctla2a
Ssc5d
Bmp4
Pf4
Ccl9
Ccr1
Igfbp2
Adgre1
Pla2g7
Mmp2
Sdc1
Coro1a
Alox5ap
Lcp1
Clec4n
Il4ra
C5ar1
Apobec1
Fcer1g
Ncf1
Fcgr3
Ctss
Ccl3
Ccl6
Trem2
Mfap2
Spi1
Unc93b1
Tyrobp
Clec4n
Il1rn
C1qc
Igf1
Col3a1
Ccr5
C1qb
C1qa
Mrc1
Ccl17
MEF
MGT+DMSO
MGT+IMAP
E
IMAP Downregulated Immune Related
Genes GO Molecular Function Analysis
Chemokine activity
Chemokine receptor binding
Receptor regulator activity
CCR chemokine receptor binding
Receptor ligand activity
C-C chemokine receptor activity
Molecular function regulator
0
2
4
6
8
10
-log p value

### Figure S3

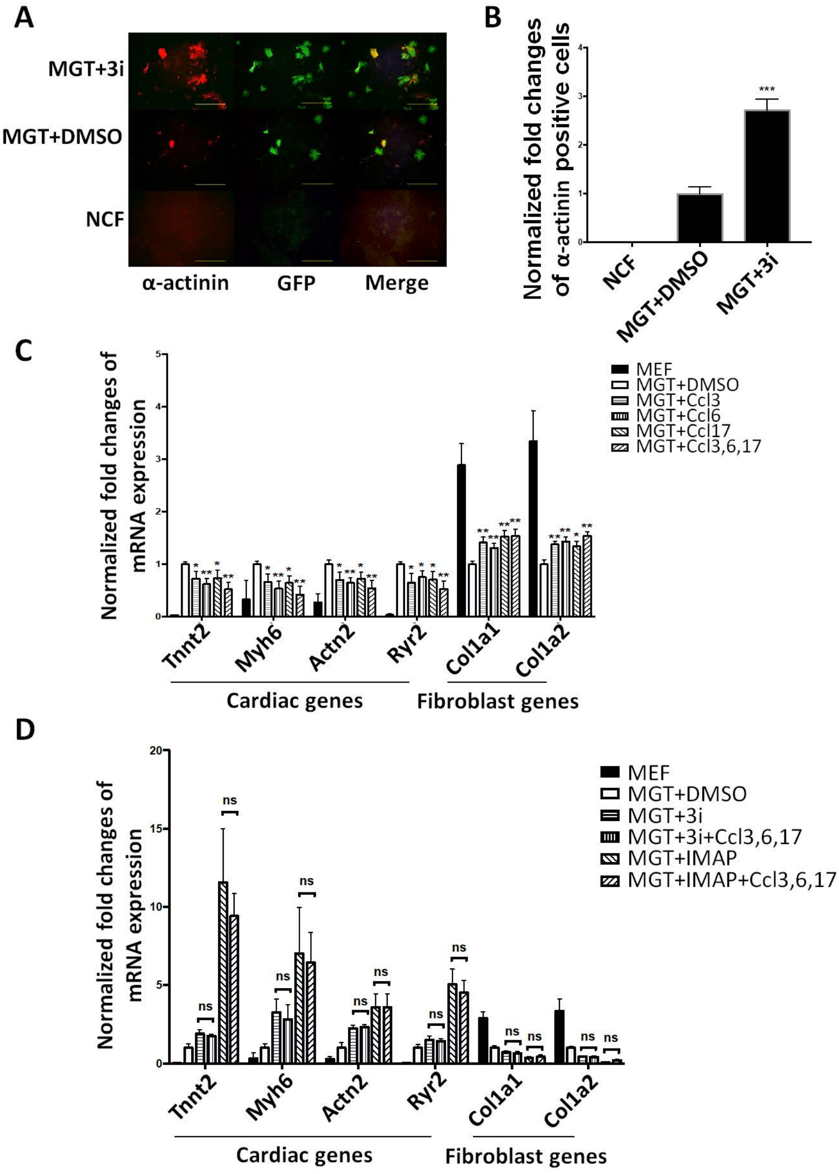
A
MGT+3i
MGT+DMSO
NCF
α-actinin
GFP
Merge
B
Normalized fold changes of α-actinin positive cells
***
NCF
MGT+DMSO
MGT+3i
C
Normalized fold changes of mRNA expression
MEF
MGT+DMSO
MGT+Ccl3
MGT+Ccl6
MGT+Ccl17
MGT+Ccl3,6,17
Tnnt2
Myh6
Actn2
Ryr2
Col1a1
Col1a2
Cardiac genes
Fibroblast genes
D
Normalized fold changes of mRNA expression
MEF
MGT+DMSO
MGT+3i
MGT+3i+Ccl3,6,17
MGT+IMAP
MGT+IMAP+Ccl3,6,17
ns
Tnnt2
Myh6
Actn2
Ryr2
Col1a1
Col1a2
Cardiac genes
Fibroblast genes
